# Genetic Distance Calculation based on Locality Sensitive Hashing

**DOI:** 10.1101/2020.04.06.027250

**Authors:** T. Pathirana, S. Bandara, G. Gamage, N. Gimhana, A. Wickramarachchi, V. Mallawaarachchi, I. Perera

## Abstract

Measuring the genetic relatedness between different species is one of the major challenges in the field of phylogenetics. Genetic distance calculation based on DNA data is highly using a mechanism to determine inter species relationships. Genetic distance computation can be further bifurcated as alignment-free sequencing and alignment based sequencing. With this research we are presenting alignment free genetic distance calculation technique which is based on locality sensitive hashing(LSH). By this approach we are hashing large DNA sequences into numeric arrays and make comparison more efficient and simplified.

## I. INTRODUCTION

Genetic distance is the degree of genetic difference (genomic difference) between species or populations which is measured by some numerical method [1]. The average number of nucleotide differences per gene is a measure of Genetic Distance. Many other molecular data can be used for measuring genetic distances. Genetic distance is an important factor in the process of reconstructing the history of the population. As a result of many kinds of research on genetic distance suggests that Saharan, African and Eurasian people diverged about 100,000 years ago[2]. Another key aspect of genetic distance is that it is used for understanding the origin of biodiversity. The different breeds of domestic animals are investigated in the purpose of determining which breeds should be protected to maintain diversity [3].

The phylogenetic tree [4] is a branching diagram or “tree” showing evolutionary relationships between different species based upon their physical and genetic characteristics. Genetic distance calculation can be considered as the main tenacity in Phylogenetic tree construction.

Genome Sequence alignment is a methodology that is currently using to identify regions of similarities and dissimilarities between sequences to calculate the genetic distances. Due to the limitations of Alignment based method Alignment Free methodology was introduced. Our approach of using LSH (locality-sensitive Hashing) tends to be used in applications related to phylogenetics without using alignment.

Calculating genetic distances between Species using LSH (Locality Sensitive Hashing) is our approach. Since DNA sequences are larger, existing LSH implementations cannot expect the maximum accuracy. Here we are introducing a new approach of creating minHash objects by partitioning sequences for comparison. Our approach tends to give convenient results which are fast and accurate.

As a primary step of Phylogenetic tree construction, the methodology we are proposing here can obtain the distance matrix by calculating inter-species distances which makes a tremendous step in the field of bioinformatics. As the next phase the distance matrix we calculated can be used to predict ancestors and cluster species into groups based on their common characteristics.

## II. BACKGROUND

Genetic distance is the way of representing similarity or dissimilarity between DNA sequences. Genome sequence alignment[5] is a method which is used to identify similar regions in sequences[6]. There are three types of genome sequence alignment types. They are global, local and glocal. Global alignment is used when the sequences have more similarities and local alignment is used when sequences have less similarities. In global alignment, it aligns every remnant in every sequence. Glocal is a combined method which attempt to find the best possible alignment of sequence.

BLAST[7] (Basic Local Alignment Search Tool) is one of the most popular alignment based tools in bioinformatics because of the functionalities and simplicity provided by the tool. It was developed by a group of scientists with the association under NIH (National Institutes of Health). In alignment free methods such as BLAST does not consider the mismatches between sequences[8]. Because of that it performance get decreased when the identity of the sequence gets increases. Another major drawback of alignment based methods is that they show less accuracy when one sequence is a repetition of another sequence. As an example, human DNA sequence is a repetition of mouse DNA sequence. But when use alignment based methods such as BLAST to evaluate similarity between those species it return lower similarity value which is incorrect. Before applying an alignment based algorithm it has to complete a considerable amount of preprocessing steps to make sequences applicable for the tool and it is time consuming[9]. This is an overhead when use alignment based approaches. Another challenge in alignment based tools is that multiple sequences cannot be processed in real time. However, most of the modern approaches of alignment use prefix indexes [10] and compression transformations such as Borrow Wheeler Transform.

Because of the imperfections in alignment based algorithms, alignment-free approach were introduced. This is a method which quantifying sequence similarity without involving alignment at any level of the algorithm[11]. Since alignment free methods has a linear complexity which depend on the length of sequence, they are computationally inexpensive. In addition to that they do not rely on assumptions regarding the evolutionary trajectories of sequence changes as alignment based methods. However alignment-free methods are focusing on occurence of sub-sequences, most of them are memory intensive. Compared to alignment based methods, these methods are still at the development level to applying in phylogenetic applications. There have more than 100 techniques [12] in alignment free category with having same characteristics we discussed above.

In current state word frequency approach is widely used to calculate genetic distances and phylogenetic applications other than alignment based approaches[13] because of their less computational power requirement than alignment based approach[14]. Use feature frequency profile(FFP) of whole genomes for comparison is one approach in alignment free method. Another alignment free approach is Comparison Vector(CV) which calculates the normalized frequency of each possible kmer of the sequence[15]. In both FFP and CV, it based on frequency of word presence in genome sequence and calculate similarity using cosine distance function [16].

But these methods do not consider the positions of sub sequence [17].

Return Time Distribution (RTD) is also an alignment free method which is different from the above-mentioned methods. It considers the amount of time taken to reappear a k-mer instead of word count [18]. As in summary these methods are creating a signature to represent a genome sequence.

In alignment free methods, the size of the signature which is used to represent the genome sequence get increased with the size of the genome sequence which is memory consumes. This is the major drawback in existing alignment free methods. As a solution for this we are proposing an approach based on Locality Sensitive Hashing (LSH).

LSH is a technique to find the near similar character sequence with high probability and efficiently [19]. This technique is most commonly used to find similar documents. LSH is deviates from conventional hashing techniques because its hash collisions are maximized, not minimized. Hashing-based approximate similarity search algorithms can be divided into two major categories. One of them is data-independent methods such as LSH. the other category is data-dependent methods such as Locality Preserving Hashing [20][21]. LSH algorithm can be subdivided into three steps. Those are shingling, minhashing and LSH. In shingling step, we convert each document into a set of characters which has constant length. Minhashing step is the most prominent step in LSH. It is a technique to estimate the similarity between two sets in very efficient manner. Originally this was designed to identify the similar web pages to disregard them from search results. In minhash step, it converts each document in to set of hash values which can be used to calculate similarity between documents. The size of the minhash object does not depend on the size of the sequence.

## III. RESEARCH MATERIALS AND DATASETS

### A. Genomic Datasets

The whole-genome sequences were downloaded from the NCBI database (ftp.ncbi.nlm.nih.gov/genomes/genbank/). For the comparisons and results we used genomes of Bacteria [22].

### B. Development Tools and Environmental Configurations

Python was the primary programming language used to implement algorithms.

Testing Environment: 8GB RAM, 4 cores of CPU, 1TB Storage, Ubuntu 16.04

## IV. METHODOLOGY

To overcome the above-mentioned drawbacks which are in the currently existing methods, we propose an approach to calculate the similarity between DNA sequences using Locality Sensitive Hashing (LSH). Since the length of a DNA sequence is higher than a normal document, currently existing LSH implementations cannot achieve the expected accuracy of DNA sequence comparing. Because in the normal LSH method, it converts a DNA sequence into a min-hash object and represents the DNA sequences using those min-hash objects. But when the DNA sequences are getting larger, the accuracy of this representation gets decreased. In our approach, we are using a DNA partitioning mechanism to overcome this drawback.

Our approach consists of four major steps. As an output of this approach, it gives the pairwise similarity between DNA sequences.

### Part A - Minhashing

Minhashing is the major step of our approach which generate a signature for a given sequence by creating a minhash object which consist of minimum hash values. Within a minhash object it contains two arrays which are called hashvalue array and permutation array. Following Algorithm 1 shows the process of initiating a minhash object by initiating those arrays. Number of minimum hash values that need to be generated for sequence is denoted by the number of permutations. The permutation array is used as a parameter for random bijective permutation function which is used to update the minimum hash values. We take a mersenne prime as the upper margin of those random value. All the hash values in hashvalue array are set to maximum hash value.

#### Algorithm 1: Initiate Minhash Object

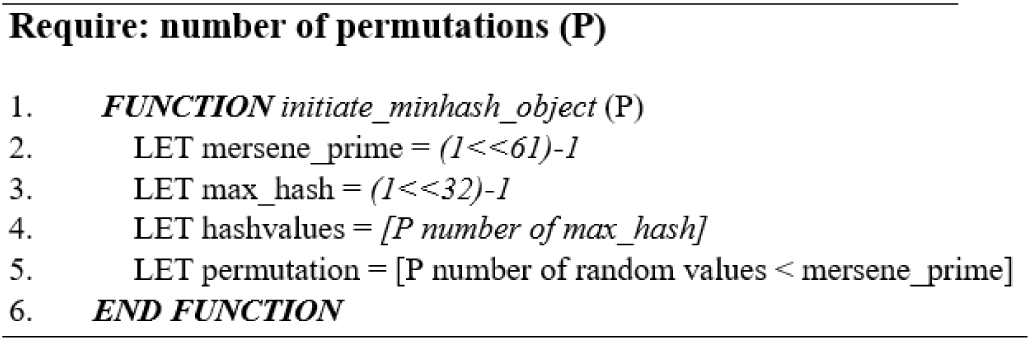

In Algorithm 2 it illustrate the procedure for minhasing a sequence. First we need to initialize the a minhash object as explained above. Then it needs to divide the DNA sequence in to constant length of subsequences which are called as “shingles”. After generating the shingles, they need to feed for an updating method to update the minhash object according to the shingles.

#### Algorithm 2: Create Miuhash for a Sequence

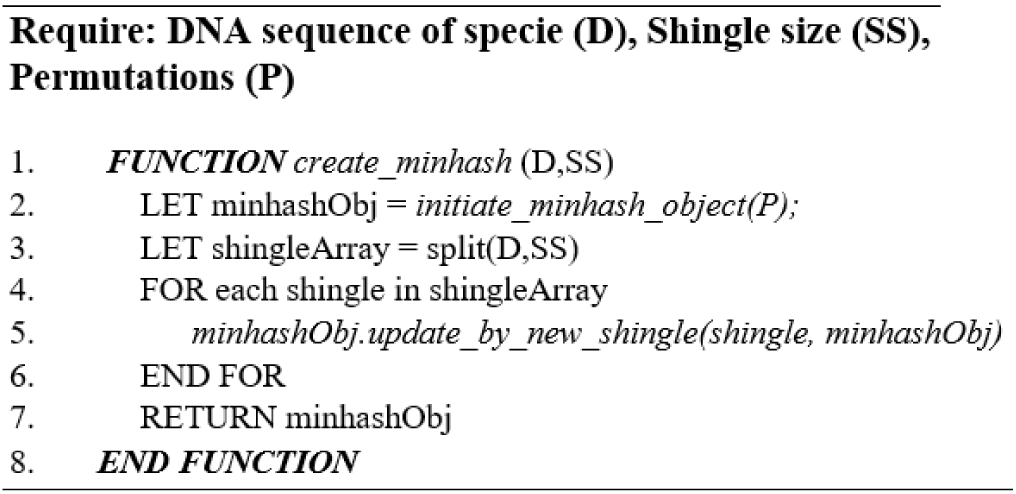

Following Algorithm 3 elucidate the updating mechanism of a minhash object according to a shingle. Target of this mechanism is to change the minimum hash values of minhash object if the fed shingle creates a hash value which is less than the existing hash values.

#### Algorithm 3: Update Miiihash Object

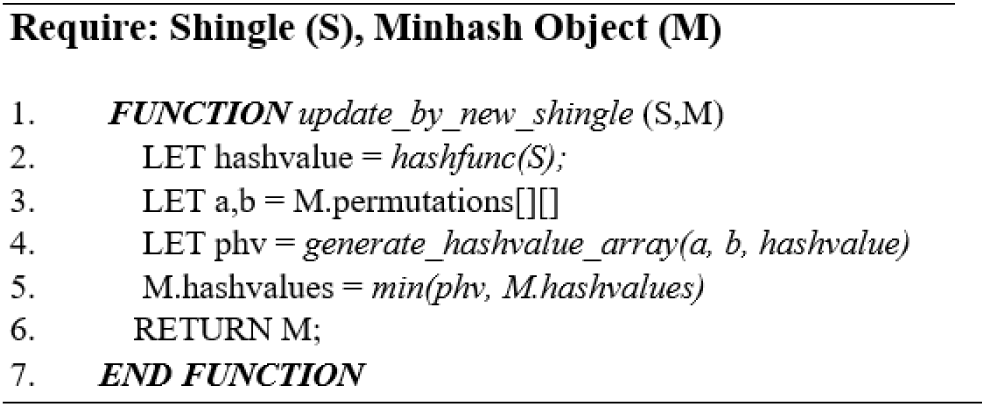

### Part B - Partitioning mechanism

In our methodology, we are representing the DNA sequences using their minhash objects. These minhash objects consist of constant number of minimum hash values of the DNA sequences. If the shingles which were forged those minimum hash values are diffused in the DNA sequence uniformly, the accuracy of the representation increased. Since the size of a DNA sequence are at high, the diffusion of the minimum hash values tends not to be uniform.

As a solution for the above drawback, we are introducing a partitioning mechanism to increase the accuracy of minhasing. Before calculating the minhash object for a given DNA sequence, we are dividing the DNA sequence into predefined number of partitions. Then each partition is minhashing by considering them as different sequences. After minhaashing all the partitions, we are getting set of minhash objects for single DNA sequence and each minhash object consists of constant number of minimum hash values. Because of the partitioning it is possible to achieve the approximate uniform diffusion of minimum hash values in the DNA sequence.

Figure 1 and figure 2 illustrates how partitioning provides the approximate uniform diffusion for minimum hash values.

**Figure 1.**
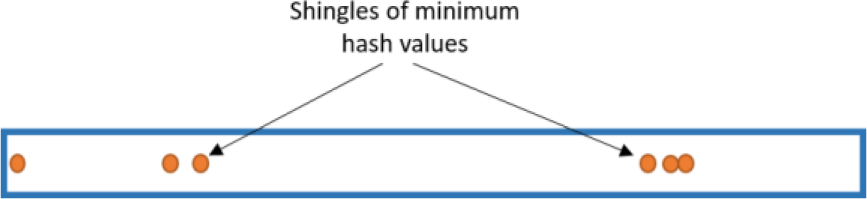
minimum hash values without partitioning DNA sequence.

**Figure 2.**
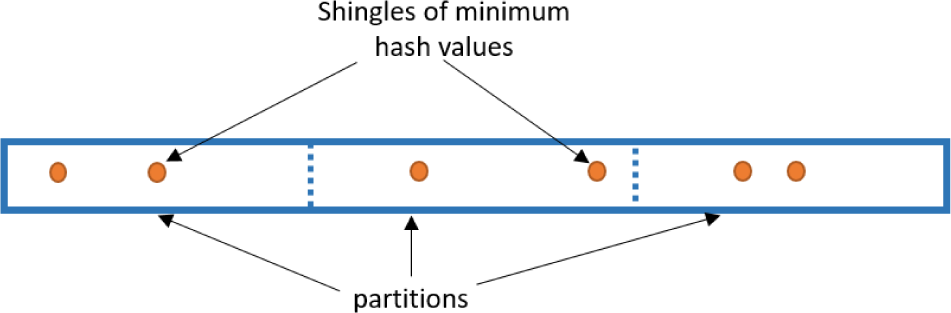
minimum hash values with partitioning DNA sequence.

In figure 1, it shows the minimum hash values are representing a lean region of the DNA sequence.In figure 2, it illustrates that when the partitioning is used, it provides the uniformness of the diffusion of the minimum hash values. And also it guarantees the uniformity of the diffusion to a certain level. The uniformity can be increased by increasing the number of partitions.

### Part C - Parallelizing Minhashing

The partitioning Approach we used above is completely independent process for each of the DNA Sequences and as we observed in the earlier stages it took much more time to Hash the Sequences. Our main target was to partition each sequence before minhashing to increase the accuracy of the representation.

Hence,as a solution for the above drawback, our next approach was to do our main task minHash running concurrently and in parallel. Here we used modules which provides interfaces for running tasks using pool of thread. After partitioning is done each thread which is in the thread pool accupie a partition of the sequence and generate a minhash object for that particular partition and store it in a global array. When all the partitions complete by generating minhash object, it combines all the minhash objects and concatenates them to create a single minhash object for the particular DNA sequence as shown in figure 3. This approach is more efficient than generating minhash objects sequentially.

**Figure 3.**
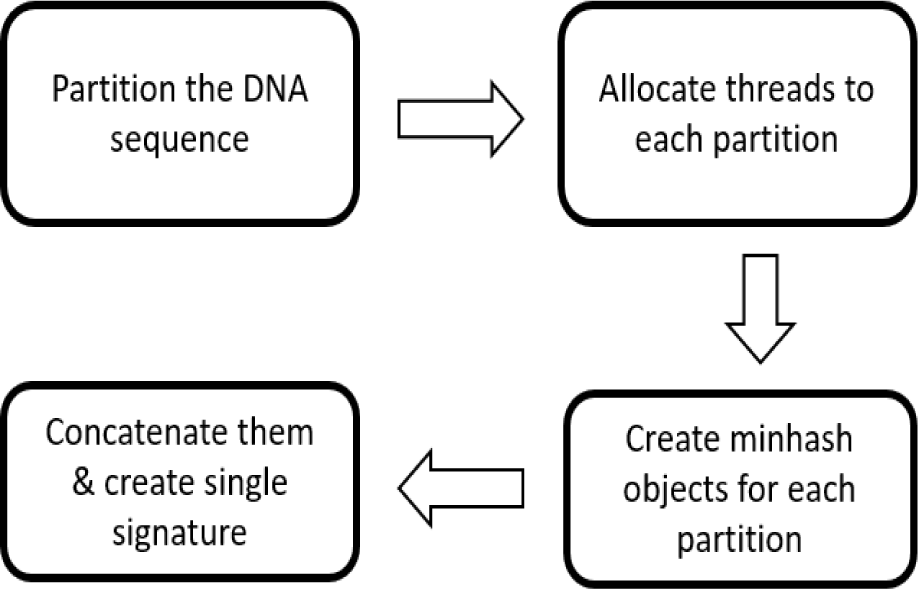
process of generating a minhash object.

### Part D - Comparing

Our Main Target was to obtain the Similarity between independent each species. As our step wise approaches we first obtain the minahash Objects which consists of Different MinHash minimum Values. When Comparing with the above two steps Comparing MinHashes and obtain the similarities takes lesser amount of time.

In this approach we obtain the Jaccard similarity[23] between the obtained DNA Minhash objects which consists of minimum hash values. The Jaccard Similarity coefficient is a statistic used for determining the similarity and diversity of sample sets. It is the coefficient where obtained by division of intersection of two samples by the union of the Sample sets as given in the following equation.

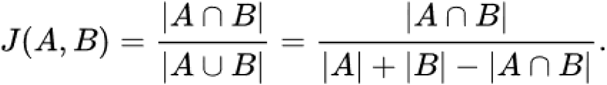

Jaccard coefficient outputs the similarity between the Minhashes obtained for different species in an efficient way.

The LSH based methodology which we partition each sequence and obtain minhash objects with minimum hash values is a novel approach that we used to compare species. Using Concurrently minhashing approach gives efficient results than the traditional approach.

## V. RESULTS AND EVALUATIONS

In this section we are mainly focusing on the evaluations done on genome Comparison using LSH. When Using LSH for our approach we had to consider some parameters which are partition time and Shingle size to obtain results with maximum accuracy of the distances between the species. When the partition size increases Comparison time decreases. Because when the partition size increases, the no of partitions decreases. Hence the partitions to compare decreases, which decreases comparison Time as shown in Figure 4.

**Figure 4.**
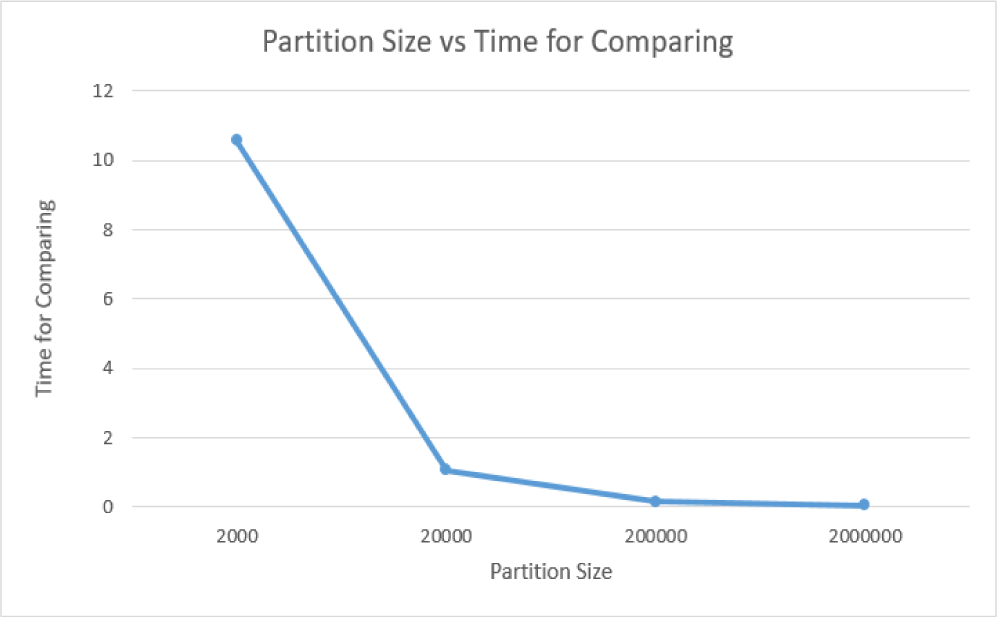
Partition size vs Time for comparing graph.

Our next approach is to obtain minHash objects from the partitions we obtained. But time taken for minHashing varies with the Partition Size we uses as shown in Figure 5. When partition size increases, time taken for minHashing increases upto some limit and then decreases.

**Figure 5.**
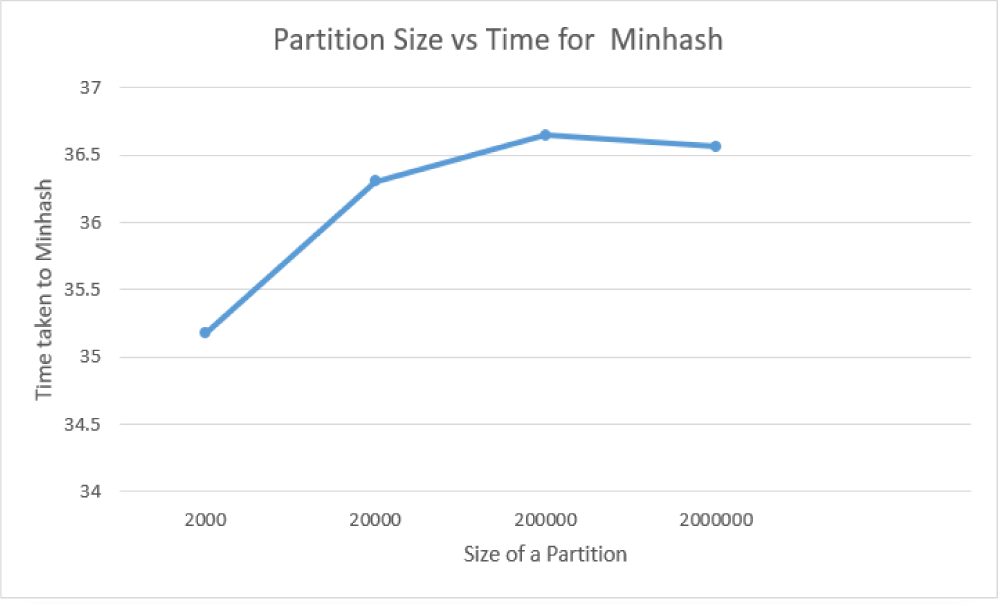
Partition Size vs Time for Minhashing graph.

For our evaluations we had to select the exact partition size which maximizes the accuracy and efficiency. By considering Figure 4 and Figure 5, we selected 20000 as the partition size for our approach.

With the shingle size we use, the minhash time varies as shown in Figure 6. We observe that the MinHash Time decreases with the Shingle Size. But when we increase the shingle size it is difficult to obtain common shingles between species. There for Accuracy decreases. Although, the minHash Time is high we use shingle size of 5 for our evaluation with the Consideration of the accuracy of comparing Species.

**Figure 6.**
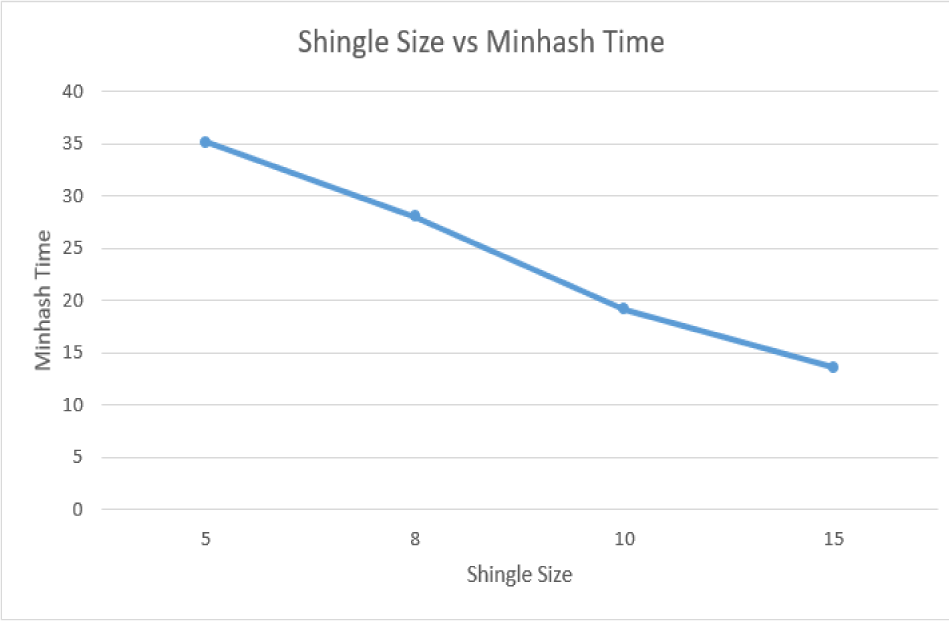
Shingle size vs Time for Minhashing graph.

## VI. DISCUSSION AND FUTURE WORK

Our focus of this paper is to compare Species using their DNA sequences based on our new approach of using LSH (Locality sensitive hashing) for genome comparison. Rather than using the traditional approach of using LSH we introduced a new method of using LSH by partitioning DNA sequences. The main important decision we had to consider with our approach is the partition size and number of shingles which gives the maximum accuracy of the distances between the species. The Final outcome of the approach that we used to generate genetic distances between species is to generate an accurate distance matrix efficiently, in consideration of obtaining phylogenetic tree. The distance matrix is playing a major role in tree construction. The reliability of the tree as well as the evolutionary relationships of taxonomies depends upon the accuracy of the distance matrix we obtained.

As our next step we suppose to construct phylogenetic trees based on the distance matrices we are building based on this methodology. For phylogenetic tree construction we are proposing novel approach which is based on machine learning. By this it is expected to extend our exiting phylogenetic tree construction pipeline.[24][25]

## VII. CONCLUSION

Locality sensitive Hashing is a methodology that can be used with whole genome sequence comparison which can be used for different genomes with diverse lengths. We obtain the similarity between species by comparing the minhash objects that we created for a given DNA sequence. Most of the DNA sequences are quite large for minhashing. We partition the DNA sequences and makes them easier for minhashing. For a sequence all the partitions were minhashed to obtain set of minhash objects. Each minhash object consists of constant number of minimum hash values. Hence memory allocated for a DNA sequence after minhashing is constant. Therefore comparing with larger DNA sequences our approach consumes the same amount of memory. Furthermore it is computationally efficient. This simplifies the genome representation and make easier for comparison. With our approach, we obtain the similarity between species by comparing pairwise to obtain jaccard similarity between them. Furthermore, we are using a concurrent approach for generating minhash objects which is time consuming. Evaluation shows that our approach guaranteed to provide the distance matrix that compares the genomes within a short period of time.

## Notes

### Competing Interest Statement

The authors have declared no competing interest.

### Summary of Updates

Added missing citations

